# Estrogen prevents age-dependent beige adipogenesis failure through NAMPT-controlled ER stress pathway

**DOI:** 10.1101/2023.08.31.555821

**Authors:** Jooman Park, Ruoci Hu, Shaolei Xiong, Yanyu Qian, Asma Sana El-Sabbagh, Meram Ibrahim, Qing Song, Gege Yan, Zhenyuan Song, Abeer M. Mahmoud, Yanlin He, Brian T. Layden, Jiwang Chen, Sang-Ging Ong, Pingwen Xu, Yuwei Jiang

**Author notes:** To whom correspondence should be addressed: Yuwei Jiang, PhD, Dept of Physiology & Biophysics University of Illinois at Chicago; Pingwen Xu, Ph.D., Address: 835 S Wolcott Ave, MC 613, Chicago, IL 60612; Sang-Ging Ong, PhD, Dept of Pharmacology & Regenerative Medicine University of Illinois at Chicago. These authors contributed equally (Jooman Park, Ruoci Hu).

## Abstract

Thermogenic beige adipocytes are recognized as potential therapeutic targets for combating metabolic diseases. However, the metabolic advantages they offer are compromised with aging. Here, we show that treating mice with estrogen (E2), a hormone that decreases with age, to mice can counteract the aging-related decline in beige adipocyte formation when subjected to cold, while concurrently enhancing energy expenditure and improving glucose tolerance. Mechanistically, we find that nicotinamide phosphoribosyltranferase (NAMPT) plays a pivotal role in facilitating the formation of E2-induced beige adipocytes, which subsequently suppresses the onset of age-related ER stress. Furthermore, we found that targeting NAMPT signaling, either genetically or pharmacologically, can restore the formation of beige adipocytes by increasing the number of perivascular adipocyte progenitor cells. Conversely, the absence of NAMPT signaling prevents this process. In conclusion, our findings shed light on the mechanisms governing the age-dependent impairment of beige adipocyte formation and underscore the E2-NAMPT controlled ER stress as a key regulator of this process.

**Highlights:**

- Estrogen restores beige adipocyte failure along with improved energy metabolism in old mice.
- Estrogen enhances the thermogenic gene program by mitigating age-induced ER stress.
- Estrogen enhances the beige adipogenesis derived from SMA+ APCs.
- Inhibiting the NAMPT signaling pathway abolishes estrogen-promoted beige adipogenesis.

## Introduction

Human life expectancy has increased rapidly (*1, 2*). However, this substantial increase in lifespan has not been accompanied by an improved quality of life for the elderly (*3, 4*). Metabolic dysfunction, which often manifests as increased fat mass and body weight, is a major precursor to age-related disabilities (*5, 6*). On the other hand, aging also raises the risk of developing metabolic diseases, such as Type 2 diabetes, cardiovascular disease, and obesity (*7–9*), accompanied with the accumulation of adipose tissue (*10, 11*). Not only does this abundance of stored fat increase the risk of premature death, but it also results in a plethora of metabolic imbalances with the advancement of aging (*12–15*).

In mammals, there are three distinct types of adipose tissue: white, brown, and beige. White adipose tissue (WAT) is characterized by its large, unilocular adipocytes which store excess energy in the form of fat depots (*16, 17*). Brown adipose tissue (BAT) functions to maintain body temperature through non-shivering thermogenesis by expressing mitochondrial uncoupling protein 1 (UCP1) (*18*). Beige adipose tissue, discovered in the last decade, is a type of inducible thermogenic fat that typically arises within the fat pads of WAT in response to cold exposure and adrenergic stimulation (*19–21*). Compared to classical brown adipocytes, beige adipocytes have emerged as a novel cellular target in the fight against obesity and diabetes, partially attributed to their considerable recruitment potential and ability to enhances energy expenditure by consuming excess calories and produce heat (*22–26*). Furthermore, this beiging capacity confers metabolic benefits including reduced blood glucose and increased insulin sensitivity (*27–31*). However, thermogenic adipose tissue activity in both mice and humans declines with aging (*32–39*). Several processes and pathways are implicated in the age-induced decline of beige fat, such as the cellular senescence of adipose progenitor cells (APCs). However, a comprehensive understanding of the molecular mechanisms underlying age-related beige adipocyte dysfunction is still largely unclear.

Estrogens play significant developmental and functional roles in many organs of male and female humans, respectively (*40, 41*). Specifically, estrogen controls whole-body metabolism through central and peripheral mechanisms by the activation of estrogen receptors (ERα and ERβ) (*42*). The loss of estrogen and knocking out ERα in mice is associated with adipose tissue redistribution, impaired glucose homeostasis, insulin resistance, and elevated type 2 diabetes risks (*42*). This is evident by the higher diabetes incidence in men and postmenopausal women versus premenopausal women (*43, 44*). Estradiol (E2; 17β-estradiol) is an estrogen steroid sex hormone derived from the cholesterol molecule, and its levels generally decline after menopause or ovariectomies (OVX) (*45*). Furthermore, E2 is implicated in the control of thermogenesis via hypothalamic AMP-activated protein kinase (AMPK) in BAT (*46*). Decreased E2 is associated with a reduction in energy expenditure and a gain in weight (*45, 47*). In addition, levels of E2 decrease with aging in both males and females (*48*). The decrease in E2 levels with aging is concurrent with a decreased function of brown and beige adipose tissue. However, the physiological role of estrogen in beige adipose tissue has not been explored with respect to its effect on the aging process.

Here, we explored that the role of estrogen in the plasticity of perivascular APCs during aging. We found that estrogen plays a pivotal role in restoring the aged-associated impairment of beige adipogenesis in perivascular APCs and in improving energy metabolism. Mechanistically, NAMPT is essential for the formation of beige adipocytes induced by E2, primarily through reducing age-related ER stress. The formation of E2-induced beige adipocytes was inhibited by either removing NAMPT or enhancing ER stress, either genetically or with drugs. Our findings provide a mechanistic understanding of how the E2- NAMPT pathway and its control over ER stress are key to understanding age-associated beige adipogenesis failure and metabolic changes.

## RESULTS

### Estrogen ameliorates aging-related defects in beige adipocyte formation

We previously reported that cold-induced beige adipocyte formation begins to decline around six months of age and becomes deficient by ’middle-age’ (i.e., twelve months old), before many detrimental aging effects on other organ function (*32, 33*). To investigate the potential role of estrogen (E2) in this aging-related decline of beige adipocyte formation, we administered either a vehicle or E2 intraperitoneally to twelve-month-old C57BL6/J mice daily for two weeks. During the second week, the mice were either exposed to cold temperatures (cold, 6°C) or kept at room temperature (RT, 23°C) (Fig. 1A). No significant difference was found in body weight between vehicle and E2 treated groups regardless of temperature (Fig. 1B). As expected, the core body temperature in mice exhibited a significant reduction upon cold exposure compared to RT (Fig. 1C). Interestingly, mice treated with E2 managed to maintain a higher core temperature during cold exposure compared to the group treated with a vehicle (Fig. 1C). Moreover, the E2 treated group showed a significant reduction in body fat content compared to the vehicle treated group when exposed to cold, but not at RT (Fig. 1D). In contrast, there were no differences in lean mass contents between vehicle and E2 treated groups regardless of temperature (fig. S1A). Consistent with body composition, E2 stimulated group mainly decreased the weight of white adipose tissue (WAT) depots (inguinal WAT (iWAT) and perigonadal WAT (gWAT)) but not the weight of other tissues such as brown adipose tissue (BAT), retroperitoneal WAT (rpWAT), muscle, kidney, spleen, pancreas, heart, and liver (fig. S1B and S1C). Furthermore, serum triglycerides were significantly lower for the E2 treated group compared to the vehicle treated group (fig. S1D). Of note, serum total cholesterol showed no difference between vehicle and E2 treated groups (fig. S1E). We also observed that there was no difference at the hepatic triglycerides and cholesterol levels between the two groups (fig. S1D and S1E). Taken together, these results indicate that E2 enhances their ability to adapt to cold environments and may exert metabolic benefits in older mice upon cold exposure.

**Fig. 1.**
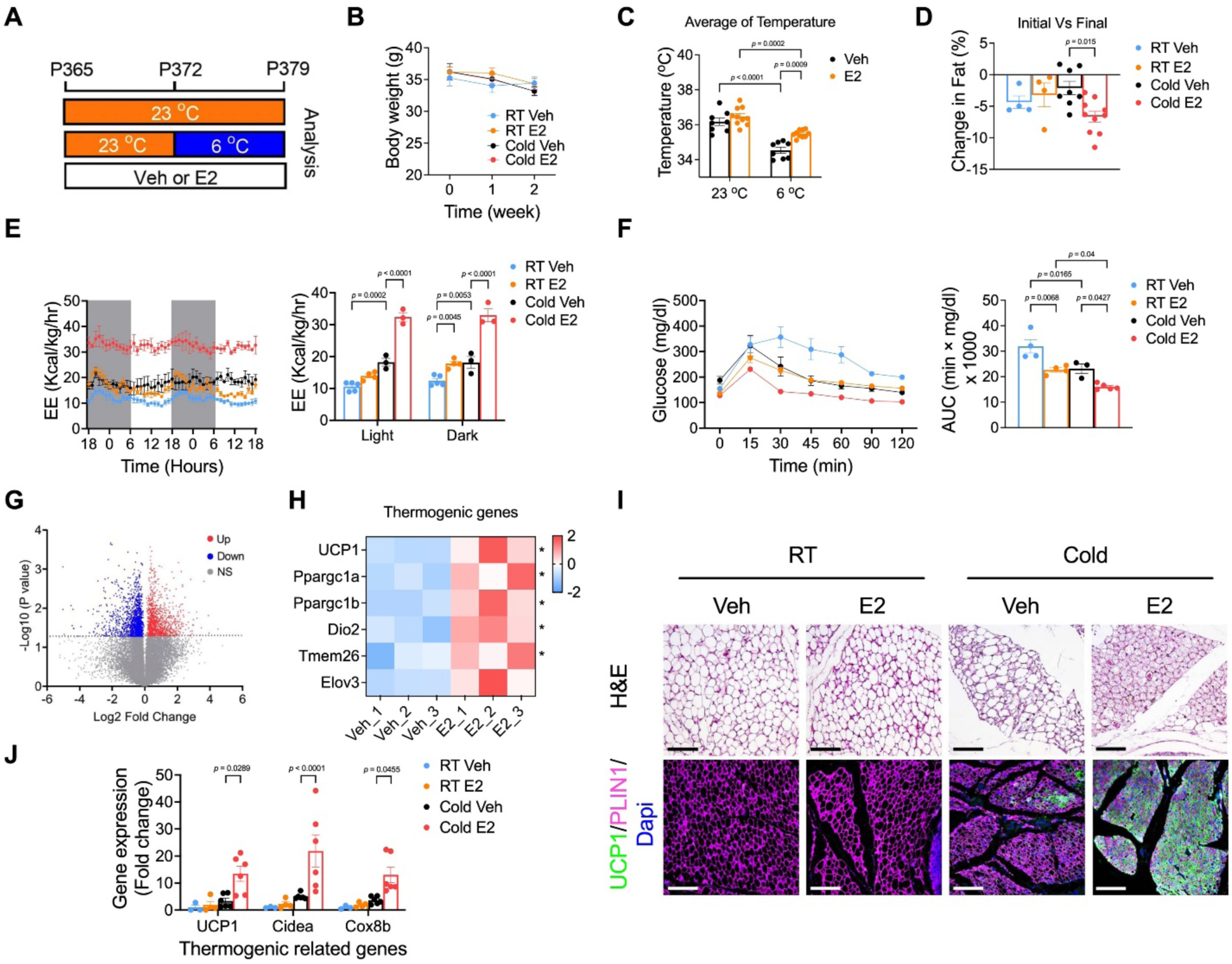
Estrogen ameliorates age-related defects in beige adipocyte formation. (A). Experimental procedure to track beige adipocytes formation in aging. 12-month-old mice were intraperitoneally injected with vehicle or E2 daily and housed in room temperature (RT; 23 ℃) for one week, followed by one week of cold exposure (6 ℃). (B-D). Body weight (B), the difference of core temperature (C), and difference of body fat composition (D). (n=10-11 per group) (E). Energy expenditure and quantification (n=3-5 per group) (F). The blood glucose levels of vehicle or E2 treated group in 12-month-old mice after intraperitoneal injection of glucose tolerance test (GTT; n=3-5 per group) and Area under curve (AUC) analysis. (G). A volcano plot of genes. Genes with log2FC ≥ 0.58 and log10P value ≥ 1.3 were considered significant. Twelve-month-old male C57BL/6 mice were divided into two groups with vehicle or E2 treated group upon cold exposure (6°C; n = 3 per group). (H). Heat map of a list of thermogenic gene expressions. (I). H&E staining and Immunofluorescence in the middle part of iWAT. Scale bar, 100 *μ*m (J). qPCR analysis of the mRNA expression of thermogenic genes in iWAT from vehicle or E2 treated upon RT (23°C) or Cold (6°C) (n= 3-6 per group). Data information: Results are presented as means ± SEM.

To further investigate the metabolic impact of estrogen *in vivo*, we measured gas exchange and whole-body energy expenditure of vehicle and E2 treated groups upon either cold exposure or RT. We found that E2 treated group showed increases in energy expenditure (Fig. 1E), oxygen consumption (fig. S1F), and carbon dioxide generation (fig. S1G) compared to vehicle treated group during both day and night cycle upon cold exposure. No significant difference was found in respiratory exchange ratio (RER) (fig. S1H), food intake (fig. S1I), and locomotor activity (fig. S1J) between vehicle and E2 treated groups under cold conditions. In addition, E2 treated group improved glucose metabolism in response to cold compared to vehicle treated group, as assessed by the glucose tolerance test (GTT) (Fig. 1F). Of note, E2 also enhanced energy expenditure and improved GTT at RT, though to a lesser extent compared to under cold conditions (Fig 1E-F). These results indicate that E2 increases energy expenditure and improves glucose homeostasis in aging mice, with these effects being markedly more pronounced under cold conditions.

To decipher the potential underlying mechanism(s) by E2 modulates energy metabolism upon cold temperatures, transcriptome profiles of iWAT in old age were investigated using RNA-sequencing (RNA-seq). We established a data threshold of ≥2 fold (log2FC ≥ 1) and identified 1,106 differentially expressed genes (DEGs) with a significant change (p-value less than 0.05) (Fig. 1G). This included 378 upregulated and 638 downregulated genes when comparing vehicle and E2 treated groups in cold-exposed older mice (Fig. 1G). Of note, the DEGs were categorized by gene ontology analysis (GO) followed by DAVID analysis (*49*). Among the genes induced by E2, we identified a clear increase in thermogenic genes (Fig. 1H). We further confirmed that E2 treated iWAT exhibited significantly higher mRNA levels of thermogenic genes induced by cold, as determined by quantitative real-time PCR (qPCR) analysis (Fig. 1J). To further assess whether E2 could rescue the failure of beige adipocyte formation, we performed Hematoxylin and Eosin (H&E) and immunofluorescence (IF) staining of UCP1 on iWAT. In line with the observed increase in energy expenditure and the expression of thermogenic genes, both H&E and UCP1 IF staining showed that mice treated with E2 possessed a significantly higher number of beige adipocytes compared to the vehicle-treated group when exposed to cold temperatures (Fig. 1I). Of note, this difference was not observed at RT (Fig. 1I). These findings suggest that E2 treatment enhances the capacity of aging mice to adapt to cold environments by promoting the iWAT beiging.

### Estrogen restores beiging potential of aged SVF cells *in vitro*

To investigate whether E2 can directly influence adipose progenitor cells for beige adipogenesis, we isolated stromal vascular fraction (SVF) cells from the iWAT of mice aged either two months or twelve months. These cells were first treated with either a vehicle or E2, then differentiated into beige adipocytes (Fig. 2A). Subsequently, they were stained with UCP1 to evaluate the number of UCP1+ beige adipocytes and analyzed mRNA levels of thermogenic genes. As expected, beige adipocytes derived from older SVF cells demonstrated a significant decrease in beiging capacity compared to those from younger SVF cells (Fig. 2B-2D). In line with our *in vivo* data, E2 markedly enhanced beige adipogenesis in old SVF cells (Fig. 2B-2D). Of note, E2 also enhanced beige adipogenesis in young SVF cells with less extent compared to old SVF cells. To further assess the functionality of E2-enhanced beige adipocytes, we evaluated the oxygen consumption rate (OCR) using seahorse extracellular flux analysis *in vitro* (*50, 51*). The basal OCR of aging SVF cells was significantly lower than that of young SVF cells (Fig. 2E-F). Consistent with increasing thermogenic gene expression levels, mitochondrial respiration was enhanced by E2 treatment in old SVF cells, but not in young SVF cells (Fig. 2E-F). Moreover, E2 stimulated both basal mitochondrial respiration and maximal mitochondrial respiratory capacity (Fig. 2E-F). Collectively, these results suggest that E2 can directly stimulate SVF cells from aging mice to differentiate into beige adipocytes and augment their function *in vitro*.

**Fig. 2.**
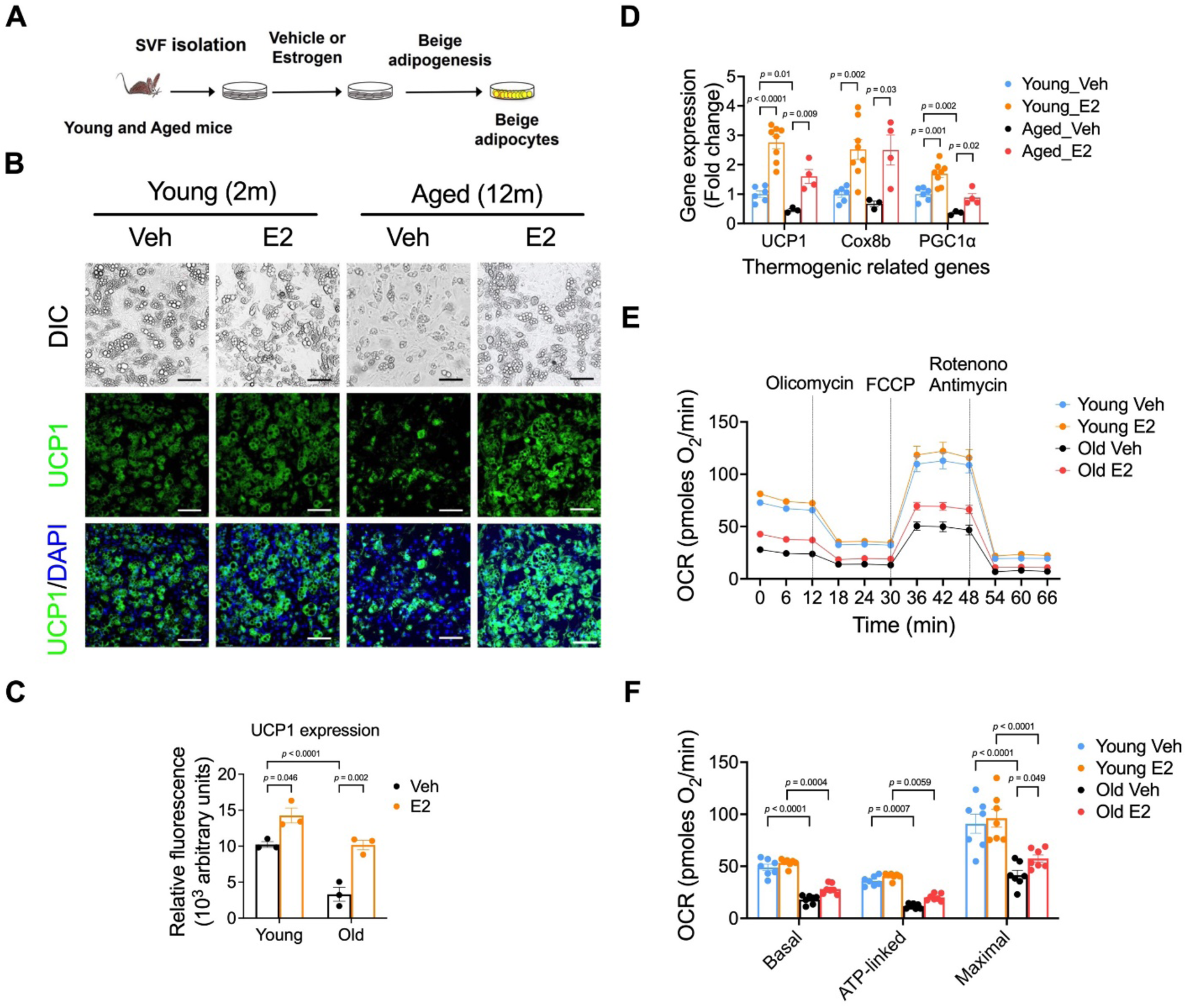
Estrogen promotes beige adipogenesis *in vitro*. (A). Schematic illustration. SVF cells isolated from iWAT of two- or twelve-month-old mice were pre-treated with either vehicle or E2 before differentiating into beige adipocytes. (B). Bright field images and immunofluorescence staining of UCP1. Scale bar, 100 μM. (C). Quantification of UCP1. (D). qPCR analysis of the thermogenic gene expression. (E). Seahorse analysis of oxygen consumption rates (OCR) in vehicle and E2 treated beige adipocytes between young (2-month) and aging (12-month). (F). Quantification of basal respiration, ATP-turnover, and maximum respiratory capacity of the samples in panel (e) (n=7 per group). Data information: Results are presented as means ± SEM.

### ER stress inducer tunicamycin attenuates cold-induced beige adipocyte formation

Our findings indicate that E2 restores the beiging ability of SVF in aging mice. To investigate the downstream pathways of the E2 effect on beiging in aging iWAT, we proceeded with GO analysis using the prior RNA-seq data (Fig 1G). Among the downregulated DEGs, “focal adhesion” and “endoplasmic reticulum (ER) lumen” GO were identified as the top cellular component pathways in cold-exposed iWAT of E2-treated mice (fig. S2A). Furthermore, Reactome analysis of downregulated DEGs revealed that ER stress-related genes, which include those suppressing beiging capacity in iWAT, were ranked at the top (fig. S2B) (*52, 53*). Therefore, we examined the expression abundance of the ER stress pathway and further validated the RNA-seq data using qPCR (Fig. 3A and 3B). Moreover, the expression levels of ER stress genes were significantly elevated in aging mice compared to young mice (Fig. 3C). In summary, our expression analysis indicate that the ER stress pathway may be one of the downstream targets that mediate E2-promoted beiging in aging mice.

**Fig. 3.**
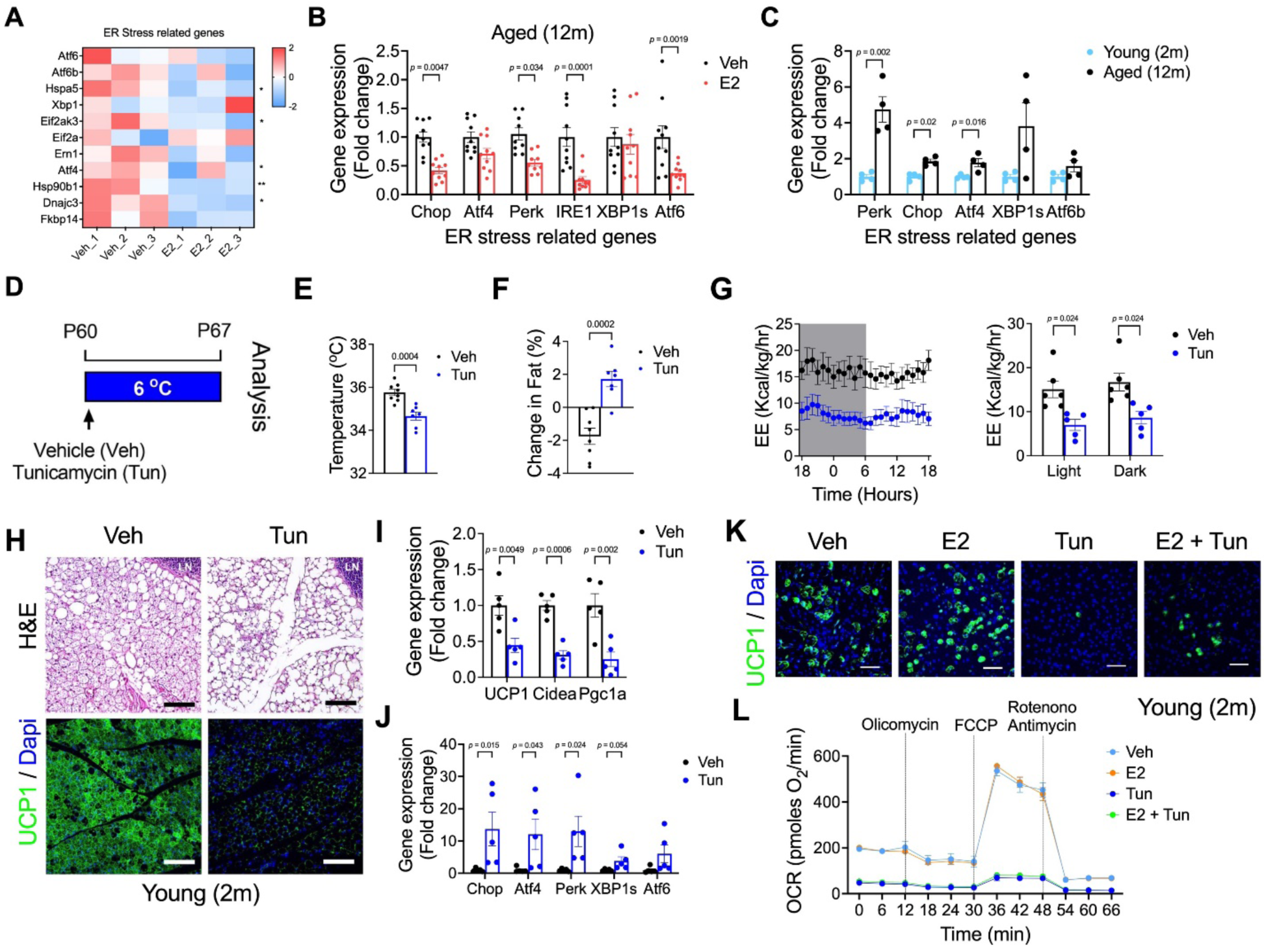
ER stress blocks beige adipocyte formation in two-month-old mice. (A). Heat map of a list of ER stress related gene expression. (B). qPCR analysis of ER stress related gene expression in vehicle or E2 treated group upon cold exposure (6°C) (n=10 per group). (C). qPCR analysis of ER stress related gene expression between young and aged (n=4 per group). (D). Experimental procedure to track beige adipocytes formation in activation status of ER stress. 2-month-old mice were intraperitoneally injected with vehicle or Tunicamycin (Tun) before cold exposure and housed in cold exposure (6℃). (E). The difference of core temperature between vehicle and Tun treated group upon cold exposure. (n=7-8 per group) (F). The difference of body fat composition between initial and final upon cold exposure. (n=7-8 per group) (G). Energy expenditure and quantification (n=5-6 per group). (H). H&E staining and Immunofluorescence of iWAT in the middle part of iWAT. Scale bar, 100 *μ*m (I-J). qPCR analysis of the thermogenic gene expression (i) and ER stress related gene expression (j) in vehicle and tunicamycin treated group (n=5 per group). (K). SVF cells isolated from iWAT of two- month-old mice were pre-treated either vehicle or E2 or tunicamycin before differentiating into beige adipocytes. immunofluorescence staining of UCP1. Scale bar, 100 μM. (L). Seahorse analysis of oxygen consumption rates (OCR) in vehicle and E2 treated beige adipocytes with/without tunicamycin. Data information: Results are presented as means ± SEM.

To investigate whether the formation of beige adipocytes was regulated by the ER stress pathway, we administered either a vehicle or tunicamycin, which is well-known to induce ER stress (*54*), to two-month-old C57BL/6 mice. The administration was carried out one dose prior to a 7-day cold exposure regimen (Fig. 3D). We found that the core body temperature in the tunicamycin-treated group was not as well-maintained as in the vehicle group (Fig. 3E). Moreover, the reduction in fat content due to cold exposure was not observed in mice treated with tunicamycin (Fig. 3F). Surprisingly, the fat percentage was found to increase, while there was no significant change observed in the lean mass content (Fig. 3F and fig. S2C). In agreement with the decrease in core temperature and increase in fat content, we observed that cold-induced beige adipocyte formation was impaired in the tunicamycin-treated group as shown by H&E and UCP1 IF staining (Fig. 3H). This beiging impairment was further confirmed by reduced thermogenesis (Fig. 3I) and induced ER stress-related genes (Fig. 3J) as detected by qPCR. Consistent with decreased thermogenesis, we found that induced ER stress by tunicamycin resulted in a significant decrease in oxygen consumption (fig. S2D), energy expenditure (Fig. 3G), and carbon dioxide generation (fig. S2E). No significant differences were observed in the RER (fig. S2F), food intake (fig. S2G), and locomotor activity (fig. S2H) between the vehicle and tunicamycin-treated groups.

To further investigate whether tunicamycin could abolish the effects mediated by E2, we isolated SVF cells from the iWAT of two-month-old mice and pre-treated them with E2 and/or tunicamycin prior to beige differentiation. In line with our *in vivo* data, the formation of beige adipocytes was inhibited in the tunicamycin-treated group, regardless of the presence of E2 (Fig. 3K). Additionally, the induction of ER stress by tunicamycin diminished both basal mitochondrial respiration and maximal mitochondrial respiratory capacity, as determined by Seahorse extracellular flux analysis (Fig. 3L and fig. S2I). Together with our *in vivo* findings, these data suggest that ER stress activation negatively regulates cold-induced thermogenesis. Furthermore, its activation can inhibit the beiging potential of both wild-type and E2-treated SVF cells.

### Perk signaling activator MK-28 prevents E2-induced beige adipocyte formation

Our findings indicate that increased ER stress activity impairs iWAT beiging. However, the ER stress signaling pathway is initiated by three ER membrane-associated sensors: activating transcription factor-6 (ATF6), double-stranded RNA-dependent protein kinase (PKR)-like eukaryotic initiation factor 2α (eIF2α) kinase (PERK), and inositol-requiring transmembrane kinase/endoribonuclease 1 (IRE1) (*55*). Thus, it remains to be determined which downstream events in the ER stress pathways are involved in beige adipocyte formation. To identify the downstream signaling, we isolated SVF cells from the iWAT of two-month-old mice and pre-treated them with a vehicle, E2, or three different ER pathway activators before differentiating them into beige adipocytes. Surprisingly, beige adipocyte capacity was decreased in MK-28 treated group, which is known activator of PERK signaling (*56*) but not the other two ER stress activators such as IXA4 or AA147, which are activators of IRE1 or ATF6, respectively (Fig. 4A and 4B) (*57, 58*). Western blotting analysis verified that there was a ∼3-fold induction of Perk phosphorylation, which is known activation form, in MK-28 treated group compared to vehicle group (Fig. 4C and 4D). These results suggest that PERK signaling may have a distinct role in E2-induced beige adipocyte formation relative to the other two ER stress signaling pathways.

**Fig. 4.**
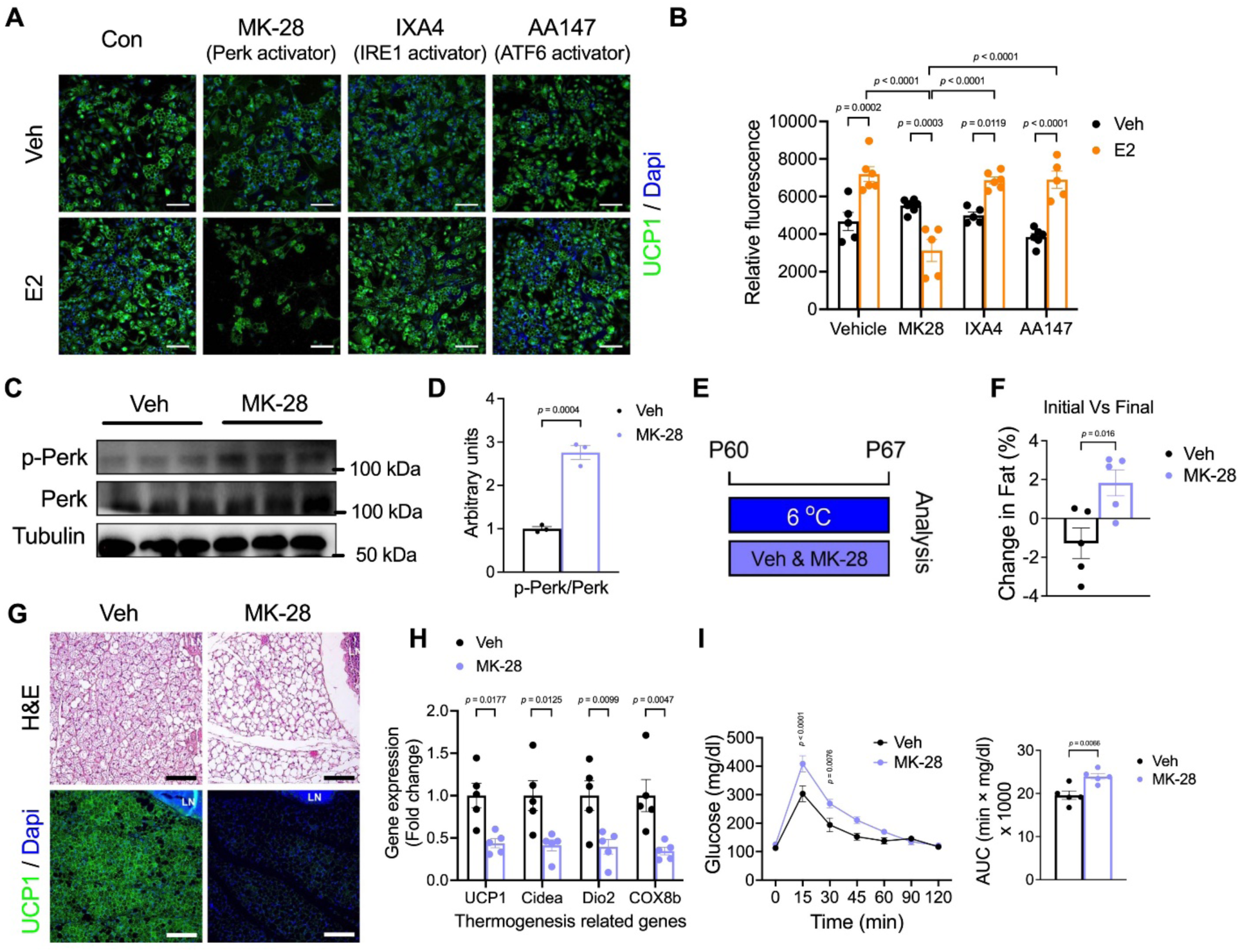
Activation of Perk signaling pathway attenuates E2-induced beige adipocyte formation. (A). Schematic illustration to track beige adipogenesis in young (two-month-old mice) and immunofluorescence staining of UCP1. SVF cells isolated from iWAT of two-month-old mice were pre-treated either vehicle, E2 or three different activators of ER stress before differentiating for beige adipocytes. Scale bar, 100 *μ* M. (B). Quantification of UCP1. (C-D). Western blot analysis of Perk and Perk phosphorylation in iWAT from vehicle and MK-28 (c), and quantification of Perk phosphorylation (d) (n=3, per group). (E). Experimental procedure to track beige adipocytes formation in activation status of ER stress. two-month-old mice were intraperitoneally injected with vehicle or MK-28 daily and housed in cold exposure (6 ℃). (F). The difference of body fat composition between initial and final upon cold exposure (n=5 per group). (G). H&E staining and Immunofluorescence of iWAT in the middle part of iWAT. Scale bar, 100 *μ*m. (H). qPCR analysis of the thermogenic gene expression (n=5 per group). (I). The blood glucose levels of vehicle or MK28 treated group in two-month-old mice after intraperitoneal injection of glucose tolerance test (GTT; n=5 per group) and Area under curve (AUC) analysis. Data information: Results are presented as means ± SEM.

To further investigate if the MK-28 treatment inhibited beige formation *in vivo*, we administrated either vehicle or MK-28 to two-month-old C57BL6/J mice for 7 days while mice were placed in cold exposure (Fig. 4E). We found that the fat content in the MK-28 treated group exhibited a significant increase compared to the vehicle-treated group (Fig. 4F). In line with the increase of fat content in the MK-28 treated group, we observed significantly reduced iWAT beiging in the MK-28 treated group compared to the vehicle group upon cold exposure based on H&E staining and UCP1 IF staining (Fig. 4G). Additionally, mRNA levels of thermogenic genes, as determined by qPCR analysis, were significantly decreased in the MK-28 treated group compared to the vehicle-treated group upon cold exposure (Fig. 4H). Next, we assessed the impact of impaired iWAT beiging on systemic glucose metabolism. Mice treated with MK-28 showed impairment in their GTT results (Fig. 4I). These data suggest that MK-28 treated PERK signaling activation disrupts beige adipocyte formation.

### SMA**+** APCs contribute to E2-induced beige adipogenesis

De novo differentiation from adipocyte progenitor cells (APCs) contribute to the formation of cold-inuced beige adipocytes (*18*). For instance, using selective genetic lineage tools, both Pdgfra+ adventitia fibroblasts and Sma+ perivascular cellular cells can emerge into beige adipocytes with the cold stimuli (*21, 59*). Our *in vitro* data thus far suggest that SVF cells directly differentiate into beige adipocytes in response to E2 treatment. To determine the specific cell populations within SVF responsible for this E2-induced beige adipogenesis, we employed two well-established APC mouse models (Sma-CreER and Pdgfrα-CreER) in combination with the permanent Rosa26R-RFP (R26RFP) reporter mice, resulting in the creation of Sma- RFP and Pdgfrα-RFP models, respectively. We administered tamoxifen (TM) to these mice at P60 and then isolated SVF cells from the iWAT of these mice for *in vitro* beige adipogenesis. These SVF cells were pre-treated with either a vehicle or E2 before beige differentiation as prior studies. Remarkably, we observed a significant increase in RFP-labeled cells arising from Sma+ APCs in the presence of E2 compared to the vehicle, but this was not observed for Pdgfrα+ APCs (Fig. 5A and 5B). In addition, the quantity of RFP- labeled UCP1+ beige adipocytes from Sma+ APCs increased upon E2 treatment, which was not observed in Pdgfrα+ APCs (Fig. 5C and 5E). As previously noted, the expression of thermogenic genes increased in the E2-stimulated group compared to the vehicle-treated group in both settings (Fig. 5D and 5F). Collectively, these results suggest that E2 effectively promotes the potential for beige adipogenesis in Sma+ APCs.

**Fig. 5.**
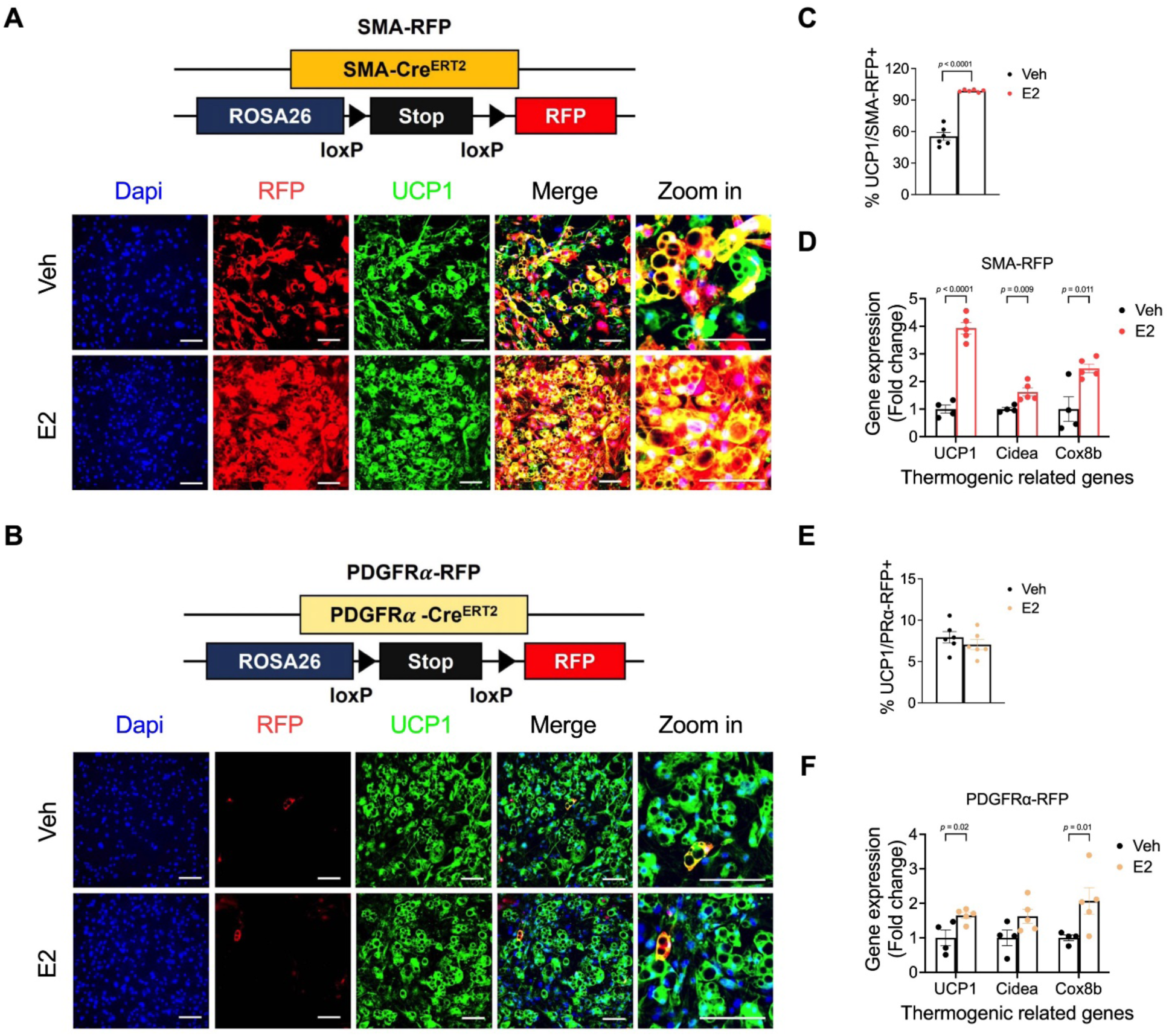
Sma+ APCs mediate E2-induced beige adipogenesis. (A). Immunofluorescence of UCP1 and RFP in Sma-RFP+ derived beige adipocytes. SVF cells isolated from iWAT of Sma-RFP two-month-old mice were pre-treated either vehicle or E2 before differentiating for beige adipocytes. Scare bar 100 *μ*M. (B). Immunofluorescence of UCP1 and RFP in Pdgfr*α*-RFP+ derived beige adipocytes. SVF cells isolated from iWAT of Pdgfr*α*-RFP two-month-old mice were pre- treated either vehicle or E2 before differentiating for beige adipocytes. Scare bar 100 *μ* M. (C). Quantification of the percentage of Sma-RFP+ cells that express endogenous UCP1 (n=6 per group). (D). qPCR analysis of the thermogenic gene expression in Sma-RFP mice model (n=4-5 per group). (E). Quantification of the percentage of Pdgfrα-RFP+ cells that express endogenous UCP1 (n=6 per group). (F). qPCR analysis of the thermogenic gene expression in Pdgfrα-RFP mice model (n=4-5 per group). Data information: Results are presented as means ± SEM.

### The loss of NAMPT in SMA+ APCs abolishes estrogen-induced beige adipocyte formation

To further identify the molecular link between ER-stress genes and E2-induced beiging, we turned back to our RNA-seq data (Fig. 1G). Intriguingly, the cellular components identified by Gene Ontology (GO) for upregulated genes included “mitochondrion,” “mitochondrial matrix,” and “peroxisome” (fig. S3A). Moreover, the molecular function of GO for upregulated genes implicated nicotinamide adenine dinucleotide (NAD+) pathways (fig. S3B). Consistent with the GO analysis, the Kyoto Encyclopedia of Genes and Genomes (KEGG) analysis revealed that the predicted functions of upregulated genes participated in various metabolic pathways and fatty acid oxidation, which is involved in mitochondrial thermogenesis (fig. S3C) (*60*). Among the upregulated genes influenced by E2, we noticed a clear increase in the expression of genes related to the NAD+ pathway, which was further validated by qPCR (Fig. 6A and 6B). Among these, we identified a notable downregulation of the enzyme nicotinamide phosphoribosyltransferase (NAMPT) - which is a rate-limiting enzyme in the NAD+ salvage pathway - associated with aging (Fig. 6B). Furthermore, the expression levels of NAMPT and associated genes were significantly decreased in aging mice compared to young mice (Fig. 6C). These findings suggest that E2 may restore the failure of beige adipocyte formation in aging individuals through NAMPT-associated NAD+ production.

**Fig. 6.**
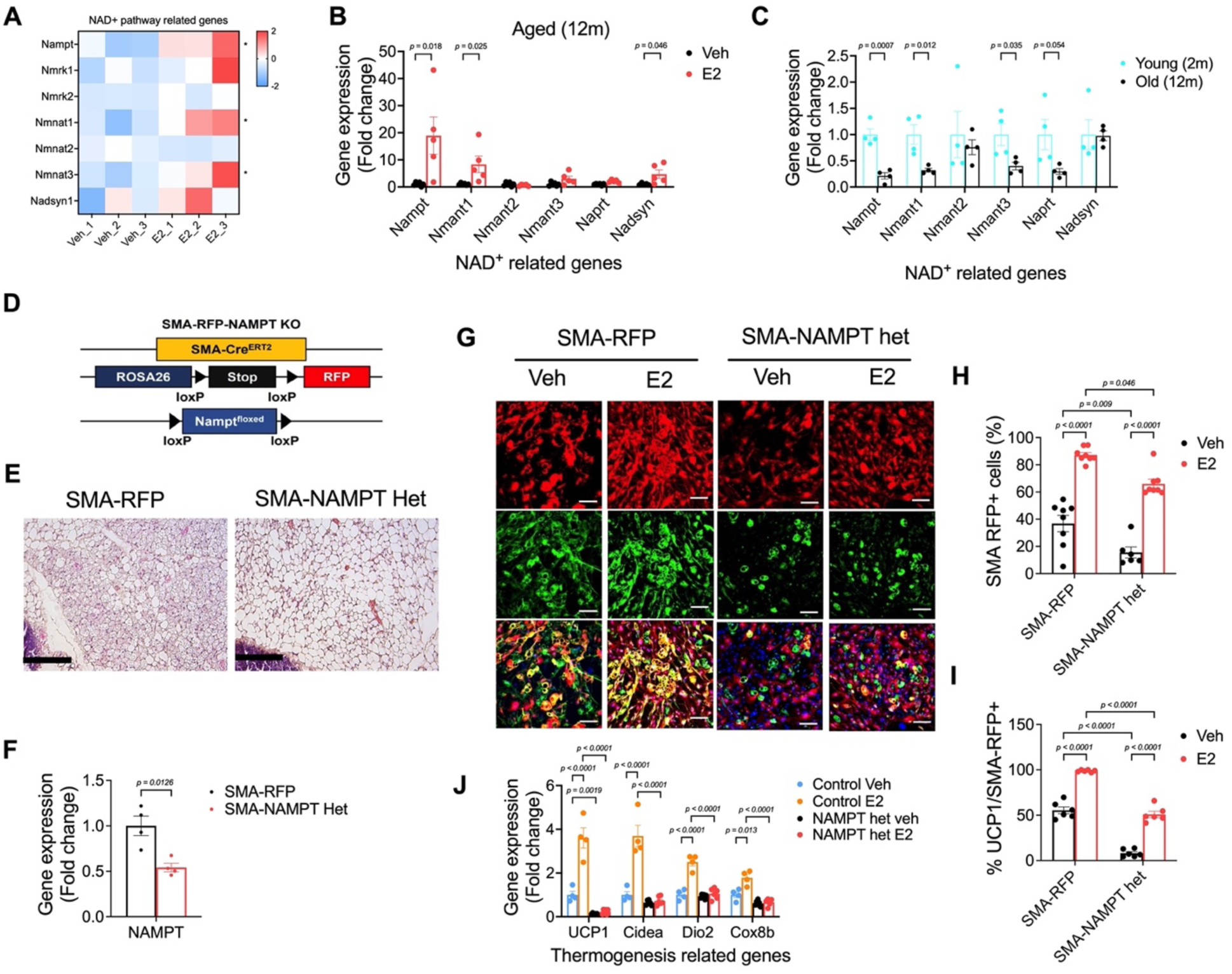
The ablation of NAMPT signaling in Sma+ APCs inhibits beige adipocyte formation. (A). Heat map of a list of NAD pathway gene expression (c). (B). qPCR analysis of NAD related gene expression in vehicle or E2 treated group upon cold exposure (6°C) (n=5 per group). (C). qPCR analysis of NAD related gene expression in between young and aging group (n=4 per group) (D). Experimental procedure to track beige adipocytes in vivo. Two-month-old conditionally knockout NAMPT in Sma+ cells (Sma-Cre^ERT2^; Rosa26R^RFP^;NAMPT^fl/fl^) were housed under room temperature after injecting tamoxifen (TAM) then, exposure cold temperature for one weeks. (E). H&E staining in control and NAMPT het mice. Scale bar, 100 *μ*m (F). qPCR analysis of NAMPT gene expression from iWAT in control and NAMPT het mice (n=4, per group) (G). SVF cells isolated from iWAT from control and NAMPT het mice in two-month-old mice were pre-treated either vehicle or E2 before differentiating for beige adipocytes. Immunofluorescence of UCP1 and RFP in Sma-RFP+ derived beige adipocytes. Scare bar 100 *μ*M. (H). Quantification of the percentage of RFP+ cells (n=6 per group). (I). Quantification of the percentage of RFP+ cells that express endogenous UCP1 (n=6 per group). (J). qPCR analysis of the thermogenic gene expression in Sma-RFP mice model (n=3 per group). Data information: Results are presented as means ± SEM.

In our above study, we identified the Sma+ APCs contributed to E2-induced beige adipocyte formation. To elucidate the role of Sma-NAMPT in mediating the effects of E2 on beige adipogenesis, we bred Sma-RFP mice with NAMPT floxed mice to create a Sma-NAMPT heterozygous (het) model (Fig 6D). At two-month-old, we administered TM to these mice, followed by testing *in vivo* beiging capabilities. In line with our hypothesis, we observed a significant decline in beige adipocyte formation in the iWAT of the Sma- NAMPT het mice (Fig. 6E). To determine whether Sma-NAMPT mediates E2’s effects on beige adipogenesis, two days post-TM injection, we harvested SVF cells from the iWAT of both control and Sma-NAMPT het mice. Prior to beige differentiation, these cells underwent pretreatment with either a vehicle or E2. A subsequent qPCR analysis confirmed a 50% reduction in NAMPT expression in the Sma- NAMPT het mice relative to the control mice (Fig. 6F). Intriguingly, irrespective of treatment type (vehicle or E2), the Sma-NAMPT het mice showed a considerable decrease in RFP-labeled cells deriving from Sma+ APCs (Fig. 6G and 6H). This reduction aligned with a decline in the number of UCP1+ beige adipocytes compared to the control mice (Fig. 6G and 6I). Furthermore, post-treatment, the Sma-NAMPT het mice exhibited fewer RFP-labeled UCP1+ beige adipocytes originating from Sma+ APCs, regardless of whether they received a vehicle or E2 (Fig. 6I). Coinciding with these observations, qPCR analyses revealed that the mRNA levels of thermogenic genes were considerably reduced in the Sma-NAMPT het group (Fig. 6J). Taken together, our findings suggest E2 stimulates Sma+ APCs to undergo beige adipocyte differentiation through a process involving NAMPT function.

### NAMPT inhibitor FK866 inhibits beige adipogenesis by inducing ER stress signaling

To further investigate the role of NAMPT in E2-mediated iWAT beiging, we administrated either E2 alone or a combination of E2 and FK866, a known NAMPT inhibitor (*61*). During the second week, the mice were exposed to cold conditions for one week (Fig. 7A). We noted no change in body weight between the two groups (Fig. 7B). However, we found E2-induced iWAT beiging was largely abolished by FK-866 co-treatment, indicated by reduced core temperature (Fig. 7D), H&E and UCP1 IF staining (Fig. 7C), and qPCR analysis of mRNA levels of thermogenic genes (Fig. 7E). To further investigate whether FK866 could inhibit the beiging effects mediated by E2, we isolated SVF cells from the iWAT of twelve-month-old mice and pre-treated them with E2 and/or FK866 prior to beige differentiation. In line with our *in vivo* data, the enhanced beige adipogenesis by E2 was largely abolished by co-treatment of FK866 assessed by UCP1 IF staining (Fig. 7F). Collectively, our findings indicate that targeting the NAMPT signaling, whether genetically or pharmacologically, could counteract the E2-mediated promotion of beige adipogenesis both *in vivo* and *in vitro*.

**Fig. 7.**
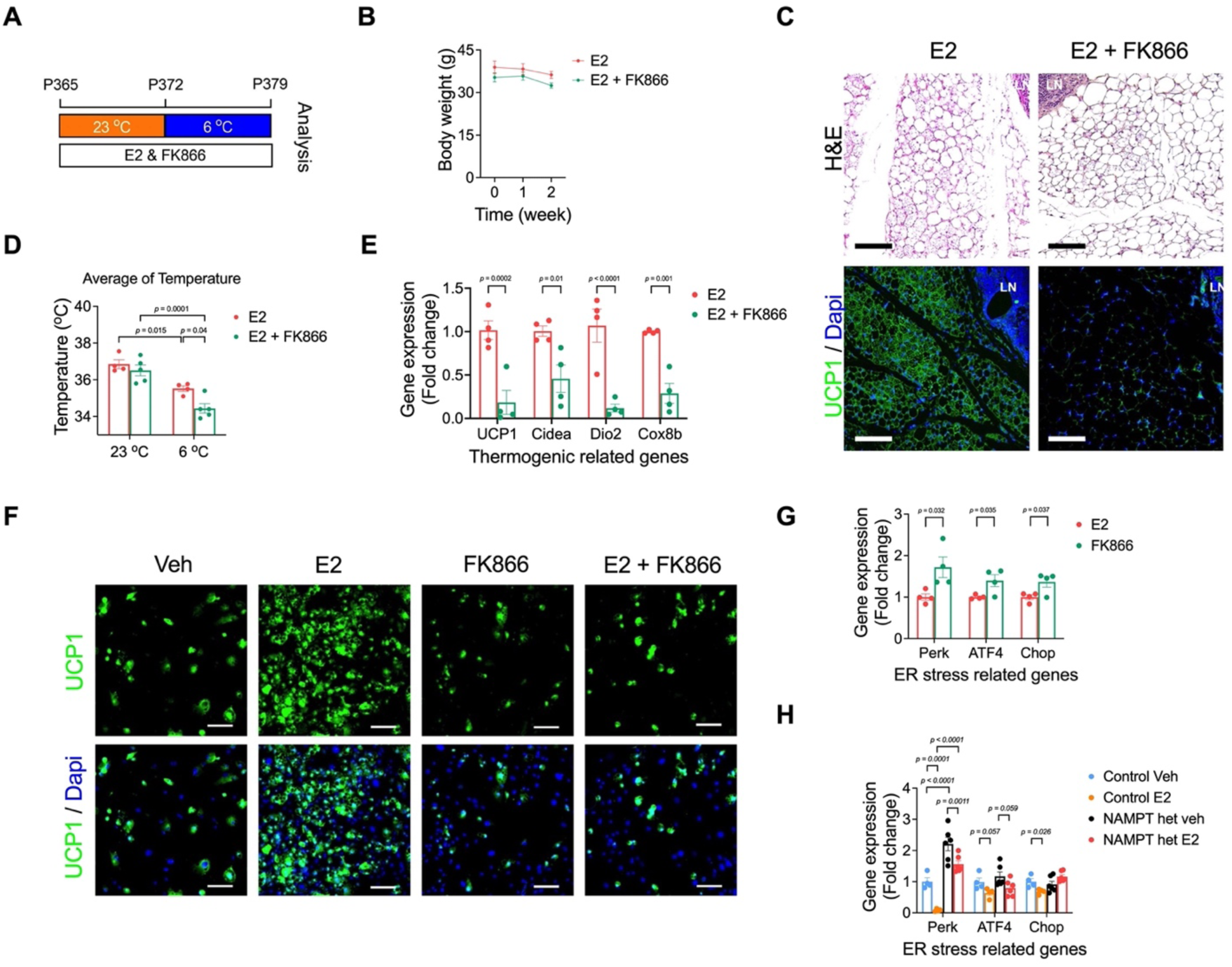
FK866 abolish E2-restored beige adipocyte formation in aging mice. (A). Experimental procedure. twelve-month-old mice were intraperitoneally injected with E2 or FK866 the other day for two weeks and housed in RT (23 ℃) for one week, followed by one week of cold exposure (6 ℃). (B). Body weight (n=4-5 per group) (C). H&E staining and Immunofluorescence in the middle part of iWAT. Scale bar, 100 *μ*m (D). The difference of core temperature (c). (n=4-5 per group) (E). qPCR analysis of the thermogenic gene expression. (F). SVF cells isolated from iWAT of twelve-month-old mice were pre-treated either vehicle or E2 or FK866 before differentiating into beige adipocytes. immunofluorescence staining of UCP1. Scale bar, 100 μM (G). qPCR analysis of ER stress in E2 or E2 comminated with FK866. (n=4, per group) (H). qPCR analysis of ER stress in control or NAMPT het mice upon vehicle or E2. (n=4, per group) Data information: Results are presented as means ± SEM.

Given that estrogen promotes the thermogenic gene program by counteracting age-induced ER stress, we subsequently investigated if NAMPT could ameliorate the beige decline of aged mice by modulating ER stress. Upon examination, we observed a pronounced induction of the ER stress pathway in the FK866- treated group relative to the vehicle-treated group (Fig. 7G). This trend was consistent with the elevated expression of the ER stress pathway observed in NAMPT heterozygous conditions (Fig. 7H). Moreover, the suppression of ER stress by E2 was compromised when one copy of NAMPT was absent, as determined by qPCR (Fig. 7H). These observations underscore the possibility that E2 could counteract the decline of beige adipocyte formation in aging subjects by inhibiting the NAMPT-driven ER stress.

### NMN administration promotes beige adipocyte formation in aging mice

NAD+ production is known to regulate key functions of numerous cellular processes, including metabolic pathways, DNA repair, and cellular senescence (*62–64*). To determine whether NAD+ is a crucial regulator of iWAT beiging in aging mice, we administered either a vehicle or nicotinamide mononucleotide (NMN), a key intermediate in NAD+ synthesis and a precursor for NAD+ production (*62*). During the second week, the mice were subjected to cold exposure for one week (Fig. 8A). We found no significant difference in body weight between the vehicle and NMN-treated groups (Fig. 8B). In parallel to the results observed with E2 treatment in aging mice, the core temperature in the NMN-treated group remained higher than in the vehicle group (Fig. 8C). In alignment with the observed increase in core temperature, we noticed a comparable enhancement of beiging responses in the NMN-treated group, as evidenced by H&E and UCP1 IF staining (Fig. 8D). Consequently, the mRNA levels of thermogenic genes were increased in the NMN-treated mice (Fig. 8E). Consistent with the upregulation of UCP1 expression by NMN in aging mice, the NMN-treated group displayed improved glucose metabolism compared to the vehicle group, as determined by the GTT (Fig. 8F). To further investigate whether NMN could rescue the beiging defects of iWAT in aged mice by modulating ER stress, we examined the associated gene expression levels. As expected, we observed a reduction in ER stress both at mRNA and protein levels in the NMN-treated group, in comparison to the vehicle-treated cohort (Fig. 8G-I). Taken together, these results indicate that NAD+ production mediated by NAMPT promotes the iWAT beiging and confers metabolic advantages similar to those of E2 in older mice exposed to cold conditions. In summary, we have identified the pivotal role of E2 signaling in counteracting the age-related decline in beige adipogenesis. This insight deepens our understanding of energy metabolism and highlights the E2/NAMPT/ER stress axis as a potential therapeutic strategy for tackling obesity (Fig. 8J).

**Fig. 8.**
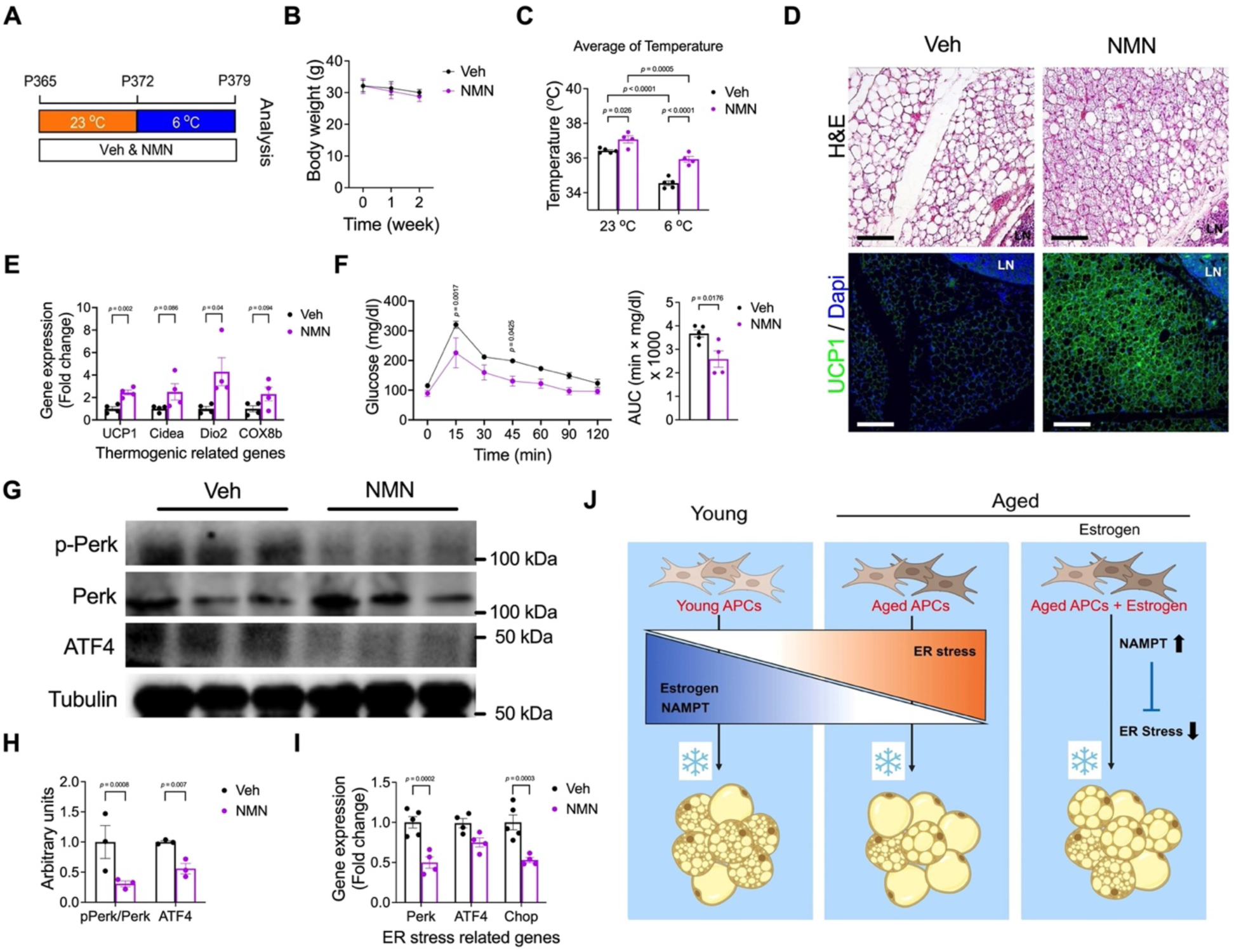
NMN administration promotes beige adipocyte formation in ageing mice. (A). Experimental procedure. One-year-old mice were intraperitoneally injected with vehicle or Nicotinamide mononucleotide (NMN) daily and housed in RT (23 ℃) for one week, followed by one week of cold exposure (6 ℃). (B-C). Body weight (B), the difference of core temperature (C) (n=4-5 per group). (D). H&E staining and Immunofluorescence in the middle part of iWAT. Scale bar, 100 *μ*m. (E). qPCR analysis of the thermogenic gene expression. (F). The blood glucose levels of vehicle or NMN treated group in twelve-month-old mice after intraperitoneal injection of GTT (n=4-5 per group) and Area under curve (AUC) analysis. (G-H). SVF cells isolated from iWAT of twelve-month-old mice were pre-treated either vehicle or E2 or NMN before differentiating into beige adipocytes. Western blot of ER stress pathway between vehicle, NMN, and FK866 in aged (G) and quantification of ER stress pathway protein (H) (n=3, per group). (I). qPCR analysis of ER stress related gene expression in vehicle or NMN treated group (n=4-5, per group). (J). Diagram illustrating the mechanisms of E2-restored beige adipogenesis in aging mice. Data information: Results are presented as means ± SEM.

## Discussions

Despite extensive research on metabolic benefits of brown/beige adipose tissue activation, age-associated beige decline has limited its therapeutic potential. In the present study, we first identified estrogen as a beiging agent, capable of reversing the age-dependent beige decline, enhancing glucose tolerance, and boosting energy expenditure in aged mice. In addition, we found that Perk signaling pathway as one of ER stress pathways, can act as downstream mediator for estrogen-induced beige formation, which consequently impair metabolic health. Moreover, our data show that estrogen can promote Sma+ APCs to differentiate into a greater number of beige adipocytes, a process that is dependent on NAMPT activity. Taken together, our studies indicate that estrogen can rescue aging-dependent beige defects through the interplay of NAMPT and ER stress pathways. As a result, estrogen can function as a potential therapeutic agent by promoting beige adipogenesis and increasing energy expenditure.

Estrogen has been known to have diverse metabolic effects including improving glucose metabolism and insulin sensitivity under different pathological conditions (*41*). The effect of estrogen and underlying mechanisms in countering age-associated beige failure have remained unexplored. In the study, estrogen is able to decrease adipose tissue content, improve glucose tolerance, and promote energy expenditure, especially under cold exposure. In addition, estrogen can exert an effect of beige adipogenesis from APCs, specifically in Sma+ cells. Previous studies have shown that beige adipocytes can be formed either through de novo differentiation or converted from mature white adipocytes (*65–67*). Our findings are in line with the notion that estrogen drives beige adipogenesis via APCs rather than converting from white adipocytes. Among the various cellular sources for beige APCs, such as Sma+ cells and Pdgfrα+ cells, our data point towards estrogen favoring beiging through Sma+ cells over Pdgfrα+ cells. However, prolonged cold exposure also recruits Pdgfrb+ APCs to differentiate into beige adipocytes; a process that we did not investigate here (*68*). It also remains to be determined which of the three known estrogen receptors - ERα, ERβ, and G protein-coupled receptor 30 (GPR30), all of which are expressed in the adipose tissue (*42*) - mediates estrogen signaling to promote beige adipogenesis within Sma+ cells. Nonetheless, our findings begin to pave the way for a novel estrogen-based cellular therapy to address the negative impacts of aging.

Endoplasmic Reticulum (ER) is a cellular organelle that is responsible for calcium storage, lipid biosynthesis, and folding newly synthesized polypeptides (*69*). ER stress occurs when disturbances in the ER lead to the accumulation of unfolded or misfolded proteins. With aging and obesity, ER stress levels rise and have been identified as contributors to metabolic syndrome (*54*). Recent research suggests a link between ER stress and the differentiation and activity of beige adipocytes. For example, the ER stress inducer, tunicamycin, has been shown to suppress Ucp1 expression in beige adipocytes and healthy young mice (*70*). Conversely, the reduction of ER stress using the agent TUDCA has been reported to stimulate beige fat formation in iWAT and counteract HFD-induced weight gain, offering potential benefits against metabolic syndrome (*71*). Our own findings align with these studies, suggesting that activation of ER stress impedes the ability of iWAT to undergo “beiging” in young mice. Furthermore, we have observed that alleviating ER stress can foster beige adipogenesis in the SVF from aged mice, as well as in the aged mice themselves. Our mechanistic investigations emphasize the pivotal role of NAMPT in regulating ER stress, which subsequently affects the differentiation and activity of beige adipocytes. However, elucidating the precise pathways connecting NAMPT and ER stress to beige adipocyte formation demands further investigation.

Therapeutically, NAD+ serves as a pivotal role in a wide range of cellular processes and energy metabolism in various organism (*72*). NAMPT, known as a rate-limiting NAD+ biosynthetic enzyme, converts nicotinamide into NMN, and then NMN adenylyltransferases will further transition NMN into NAD+ (*64*). The association between NAMPT dysfunction and metabolic impairment has been well documented. Several studies have described that adipocyte specific NAMPT deletion can cause adipose tissue dysfunction and insulin resistance in multiple metabolic organs (*73, 74*). Moreover, brown adipocyte specific NAMPT KO has been regulated energy metabolism via reducing thermogenesis in BAT (*75*). However, the role of NAMPT signaling in APC function and maintenance remains unexplored. The data we collected has shown that Sma+ cells with NAMPT deletion show beige adipogenesis defects irrespective of estrogen treatment, indicating NAMPT is involved in estrogen-mediated beige adipocyte formation. In addition, pharmacological supplementation with NMN can rescue aging dependent beige failure in old mice. It remains further investigation that whether NAMPT overexpression specifically in Sma+ cells can restore aging-dependent beige adipogenesis failure.

In summary, this work uncovers the pivotal role of the E2-NAMPT controlled ER stress pathway in regulating beige adipogenesis. Through both genetic and pharmacological manipulation of this pathway, we effectively countered the age-related reduction of beige adipocytes in aging mice, thereby promoting metabolic health. Given the benefits of beige adipocytes in fighting obesity and related issues, our findings suggest that E2-NAMPT controlled ER stress pathway could be used to boost beige adipogenesis, hoping to treatments for aging and related health problems.

## Material and Methods

### Animal

All animal experiments were performed according to procedures reviewed and approved by the Institutional Animal Care and Use Committee of the University of Illinois at Chicago under the auspices of protocol number 2021-0112. Mice were housed in a temperature/humidity-controlled environment (23° ± 3°C/70 ± 10%), and a 14:10 light:dark cycle with a standard rodent chow diet and water unless otherwise indicated. All mice were purchased from Jackson Laboratory. All animal experiments were performed on male at denoted ages per experiment. All experiments were performed on 3 or more mice per cohort and performed at least twice. Animals were euthanized by carbon dioxide asphyxiation and cervical dislocation was performed as a secondary euthanasia procedure. For fate mapping, R26R^RFP^ were purchased from JAX (#007914). Sma-Cre^ERT2^ mice were generously provided by Dr. Pierre Chambon. Drs. Sean Morrison and Bill Richardson generously provided the Pdgfrα-Cre^ERT2^ mice. For deleting NAMPT, NAMPT floxed mice were generously provided by Dr, Shin-ichiro Imai (Washington University). We crossed Sma-Cre^ERT2^ mice with NAMPT floxed and R26R^RFP^ mice. Cre recombination was induced by administering one dose of tamoxifen (Cayman, Ann Arbor, MI) dissolved in sunflower oil (Sigma-Aldrich, St. Louis, MO) for two consecutive days (50mg/Kg intraperitoneal injection).

### Pharmacological administration

Twelve-month-old male C57BL6/J mice were administrated intraperitoneally (IP) with 17β-estradiol (E2; 1µg/Kg, Cayman, Ann Arbor, MI) dissolved in sesame oil (Sigma-Aldrich, St. Louis, MO) or nicotinamide mononucleotide (NMN; 50mg/Kg, Ambeed, Arlington Hts, IL) dissolved in 1X PBS for 2 consecutive weeks. FK866 (10mg/Kg, Ambeed) was administrated into 1-year old male C57BL6/J mice for every other day at same dose of vehicle (1X PBS). To induce ER stress, 2-month-old male C57BL6/J mice were administrated one dose of tunicamycin (0.5µg/Kg, Sigma-Aldrich) or seven doses of MK-28 (1mg/Kg, Medchem Express, Monmouth Junction, NJ) via IP injection. Subsequently, the mice were subjected to cold temperature challenges.

### Metabolic phenotyping experiments

Mice were housed individually and acclimatized to the metabolic chambers (Promethion System, Sable System International, Las Vegas, NV) at the UIC Biologic Resources Laboratory for 2 days before data collection was initiated. For the subsequent 3 days, Food intake, oxygen consumption, carbon dioxide production, energy expenditure, and physical activity were monitored over a 12 h light/dark cycle with food provided ad libitum. The energy expenditure was normalized to lean body mass. The RER was calculated using the VCO2/VO2 ratio from the gas exchange data. For energy expenditure analysis, embedded ANCOVA tools in a web application CalR (*76*) were used to perform regression-based indirect calorimetric analysis. For measuring core body temperature, the probe was lubricated with glycerol and inserted 1.27 centimeters (0.5 inch), and the temperature was recorded when stabilized at the indicated time points. Body composition was measured using a Bruker Minispec 10 whole body composition analyzer (Bruker, Billerica, MA) at the UIC Biologic Resources Laboratory. For cold exposure experiments, mice were placed in a 6°C environmental chamber (Environmental & Temperature Solutions, Hoffman Estates, IL) for 7 days. Body temperature was measured using a rectal probe (Physitemp, Clifton, NJ). Blood glucose levels were measured using automated glucose meter (Contour Next, Bayer, Bayer AG, Leverkusen, Germany). Glucose tolerance test (GTT) were performed as described previously (*77*). Briefly, intraperitoneal GTT was performed after 4 hours of fasting. An injection of glucose (1.5 g/kg body weight) was given to the mice, and blood glucose levels were measured subsequently at 0, 15, 30, 45, 60, 90, and 120 min.

### Serum analysis

Blood was collected from the heart and then centrifuged at 5,000 g for 10 min. Then, the plasma was extracted, and aliquots were stored at -80°C until further analysis. The concentrations of triglyceride (TG) and cholesterol in the serum were determined using Triglyceride E test kits (Thermo Fisher Scientific, Waltham, MA) and cholesterol LiquiColor kits (Stanbio, Boerne, TX) respectively. All assays were performed according to the manufacturer’s instructions.

### Isolation of stromal vascular fraction (SVF)

SVF cells were isolated as previously described (*21*). Briefly, iWAT were dissected and digested at 37°C for 40 min in isolation buffer (100 mM HEPES, 0.12 M NaCl, 50 mM KCl, 5 mM D-glucose, 1.5% BSA, 1 mM CaCl2, pH 7.3) containing 1 mg/ml collagenase type I (Worthington Biochemical, Lakewood, NJ). After removing the floating layer containing mature adipocytes, the aqueous phase was centrifuged at 1000× g for 10 min. The pellet, including a crude SVF, was resuspended in red blood cell (RBC) lysis buffer (155 mM NH4Cl, 10 mM KHCO3, 0.1 mM EDTA) for 5 min and centrifuged at 1000× g for 5 min. The pellet was washed once with 1X PBS, resuspended, and strained through 70 mm mesh. The pallets were resuspended and seeded.

### Cell culture

The isolated SVF cells were cultured in DMEM/F12 media (Sigma-Aldrich) supplemented with 10% FBS (Sigma-Aldrich) and 1% Penicillin/Streptomycin (Gibco, Waltham, MA) at 37°C in a 5% CO2 humidified incubator. For beige adipocyte differentiation, post-confluent cells were induced by DMEM/F12 containing 10% FBS, 0.5 mM isobutyl-methylxanthine (Sigma-Aldrich), 1 µg/mL insulin (Sigma-Aldrich), 1 µM dexamethasone (Sigma-Aldrich), 60 µM indomethacin (Sigma-Aldrich), 1 nM triiodo-L-thyronine (Sigma-Aldrich), and 1 µM rosiglitazone (Sigma-Aldrich) for first 2 days and with DMEM/F12 containing 10% FBS and 1 mg/mL insulin, 1 nM triiodo-L-thyronine, and 1 mM rosiglitazone every 2 days. Fresh medium was replaced every 2 days until ready for harvest. For the pharmacological treatments, cells were pretreated with E2 (10 nM), NMN (50µM), Tunicamycin (0.5µg/ml), or MK-28 (10µM) two days prior to initiating beige differentiation. FK866 (300nM) was administered during the differentiation process.

### Histological staining

Hematoxylin and eosin (H&E) staining was conducted on paraffin sections using standard methods as described previously (*78*). Briefly, adipose tissues were fixed in formalin overnight, dehydrated, embedded in paraffin, and sectioned with a microtome at 5 μm thicknesses. For immunofluorescence staining, paraffin sections were incubated with antigen retrieval buffer (1X Sodium Citrate Buffer pH 6.0 in PBS) after 30 min at room temperature (23°C), with primary antibody at 4°C overnight, and with secondary antibody for 2 h at room temperature. Antibodies used for immunostaining were rabbit anti-UCP1 (1:400, PA1-24894, Thermo Fisher Scientific), goat anti-Perilipin (1:400, ab61682, Abcam, Cambridge, UK), DAPI (SP-8500, Vector Laboratories, Newark, CA). Secondary antibodies were purchased from Jackson ImmunoResearch. All secondary antibodies were used at a 1:400 dilution. Immunostaining images were taken using a Leica DMi8 microscope (Leica, Wetzlar, Germany) or confocal microscope (LSM 880, Carl Zeiss, Oberkogen, Germany).

### RNA-sequencing and Quantitative real-time PCR (qPCR)

Total RNA from tissue and cells was extracted using Tripure Isolation Reagent (Sigma-Aldrich) using Bullet Blender Homogenizer (Next Advance, Troy, NY) according to the manufacturer’s protocol. RNA quality evaluation (yield, purity, and integrity), cDNA library construction and illumine sequencing were conducted by Novogene (Beijing, China) as described previously (*79*). The ClusterProfiler R package was used to test the statistical enrichment of differentially expressed genes in KEGG pathways. Genes with p-value < 0.05 and log2(Foldchange) ≥ I1I were considered significantly. cDNA was generated from 1 μg of total RNA using a High-Capacity cDNA Reverse Transcription Kit (Thermo Fisher Scientific). qPCR was performed using a ViiA7 system (Applied Biosystems, Forster City, CA, USA) according to the manufacturer’s protocol. Data were analyzed using the comparative Ct method and the qPCR value were normalized by 18s rRNA expression. mRNA levels were expressed as the fold-increase relative to basal transcription levels. Primer sequences are available in Table S1.

### Oxygen Consumption rate (OCR)

Cellular oxygen consumption rate (OCR) of beige adipocytes was determined using a Seahorse XF96 Extracellular Flux Analyzer according to the manufacturer’s instructions. Briefly, SVF cells from iWATs of 2-month-old and 1-year-old mice were seeded on an XF96 plate and induced to undergo beige differentiation. The plate was then placed in a temperature-controlled (37°C) extracellular analyzer at the indicated time points. After measuring initial oxygen consumption rate (OCR), 4 *μ*M oligomycin, 2 *μ*M FCCP and 1 *μ* M rotenone/actinomycin A were then sequentially added into the plate by automatic pneumatic injection. Data were analyzed by Seahorse Wave software. Basal respiration was calculated as [OCR^initial^ - OCR^R&A^]. The ATP-linked respiratory was computed as [OCR^FCCP^- OCR^oligomycin^]. The maximum respiration rate was computed as [OCR^FCCP^ - OCR^R&A^].

### Western blotting analysis

Western blotting analysis was performed as described previously (*77*). Briefly, total proteins from tissue and SVF cells were lysed with RIPA buffer (Boston BioProducts, Boston, MA) supplemented with protease inhibitor cocktail (MedChem Express, Monmouth Junction, NJ) and then cleared by centrifugation at 21,000 g for 15 min at 4°C. The amounts of proteins were detected by a BCA assay kit (Thermo Fisher Scientific). Equal amounts of protein extracts were subjected to electrophoresis in 8–12% SDS-polyacrylamide gels and then transferred to PVDF membranes (EMD Millipore, Burlintgon, MA). The membranes were blocked with 5% non-fat milk at room temperature and then incubated with relevant primary antibodies at 4°C overnight, followed by horseradish peroxidase (HRP)-conjugated secondary antibody at room temperature for 2 h. The signals were detected by the addition of Western ECL Substrate (Thermo Fisher Scientific).

### Statistics and Reproducibility

All data are presented as mean ± SEM. Two-tailed unpaired Student’s t test (for comparison of two groups) or one-way ANOVA followed by Tukey’s test (for comparison of three or more groups) were conducted using GraphPad Prism software. All experiments were repeated two or three times with representative data or images shown. p < 0.05 was considered statistically significant in all the experiments.

### Data availability

The data that support the findings of this study are available in the methods and supplementary material of this article. Source data are provided with this paper. Additional data is available upon request from the corresponding author.

## Acknowledgments

We thank Dr. Shin-ichiro Imai for kindly providing the NAMPT floxed mouse strain. We thank members in the Y.J. laboratory for mouse genotyping and technical help. We thank Cynthia Rose Adams and Jeanette Purcell for assistance with mouse husbandry, Stefan J. Green and the Research Resources Center for RT-PCR analysis, Metabolic Phenotyping Core for analytical and phenotypical mouse measurements, and members of the Y.J. laboratory for helpful comments on the manuscript. This work was supported by grants to Y.J. from the National Institute of Diabetes and Digestive and Kidney Disease grant (NIDDK) K01 DK11177, R03 DK127149, R01 DK132398 and Pilot & Feasibility Diabetes Research & Training Center (DRTC) Award (P30DK020595).

## Author contributions

Y.J. and J.P. conceived and designed the experiments. J.P. and R.H. conducted most of the experiments. R.H., J.P., and Y.J. interpreted the experiments, wrote, and revised the manuscript, and S.X, Y.Q, G.Y., S.O., Q.S., Z.S., B. L, J. C, Y.H, A.M., and P.X. reviewed it.

## Competing interests

The authors declare no conflict of interests.

## Notes

### Competing Interest Statement

The authors have declared no competing interest.

