## Supplemental files for "Estrogen prevents age-dependent beige adipogenesis failure through NAMPT-controlled ER stress pathway"

Park *et al.*

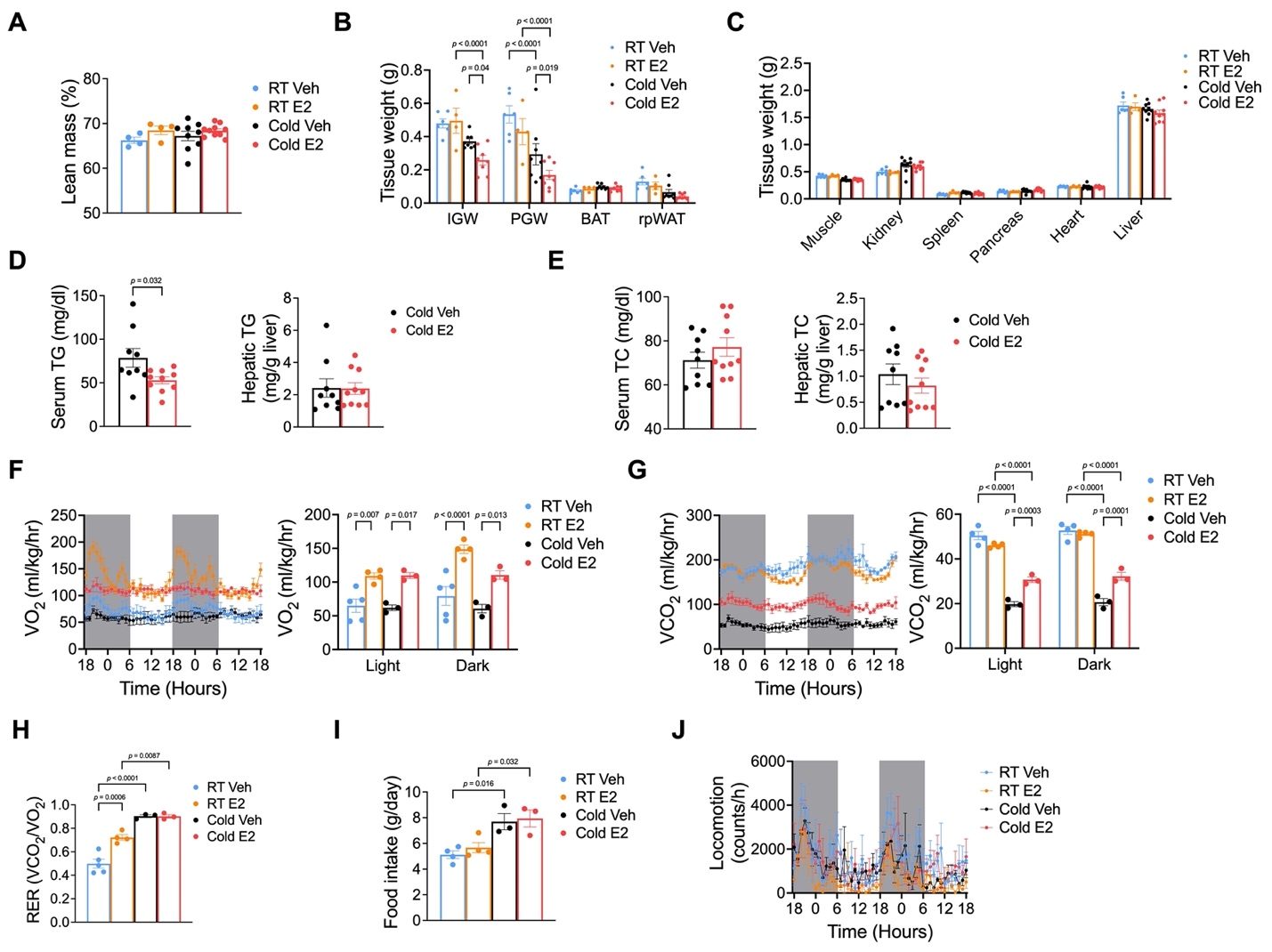

**Fig. S1. Estrogen ameliorates some age-associated metabolic phenotype impairments.**

(A). body lean mass composition (n=10-11 per group) (B-C). Weight of adipose tissues (B), and other tissues (C). (n=10-11 per group) (D-E). Serum and hepatic triglyceride (D), and serum and hepatic cholesterol (E). (n=9-10 per group) (F-J). Oxygen consumption and quantification (F), Carbone dioxygen consumption and quantification (G), RER (H), food intake (I), and physical activity (J). (n=3-5 per group) Data information: Results are presented as means ± SEM.

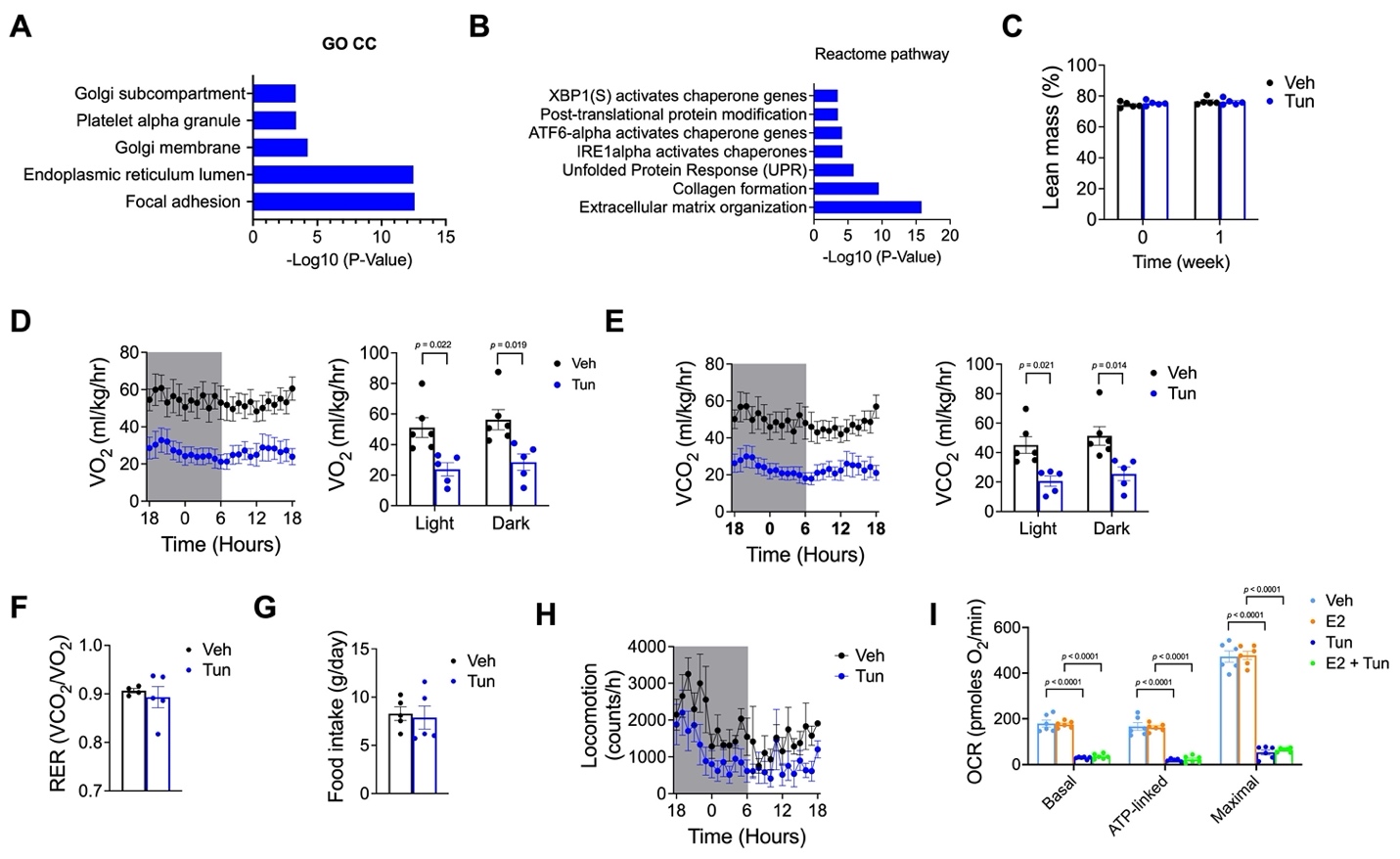

**Fig. S2. ER stress inducer tunicamycin impairs energy expenditure in young mice.**

(A). Downregulated gene ontology (GO) cellular component to E2. (B). Downregulated GO reactome pathway to E2. (C). The lean mass composition in vehicle and tunicamycin treated group. (n=5 per group) (D-H). Oxygen consumption and quantification (D), Carbone dioxygen consumption and quantification (E), RER (F), food intake (G), and physical activity (H). (n=4-6 per group) (I). Quantification of basal respiration, ATP-turnover, and maximum respiratory capacity of the samples in panel. (n=6 per group) Data information: Results are presented as means ± SEM.

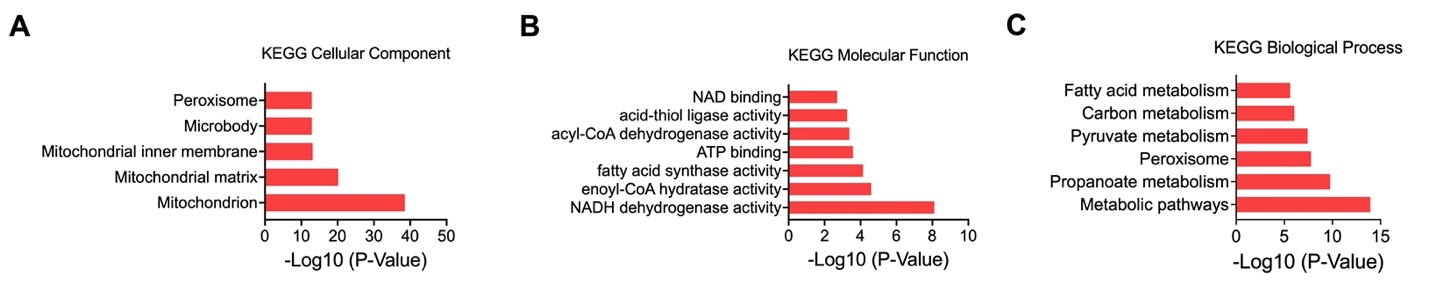

**Fig. S3. NAD**$\boldsymbol{+}$ **pathway analysis in E2-treated aged iWAT.**

(A-C). Upregulated gene ontology (GO) cellular component to E2 (A), upregulated gene ontology (GO) molecular function to E2 (B), and heat map of a list of NAD pathway gene expression (C). (n=3, per group)

**Supplementary Table 1. The primer sequences for qPCR in this Study**

| Genes |  | Sequence (5’-3’) |
| --- | --- | --- |
| 18s | Forward | GTAACCCGTTGAACCCCATT |
|  | Reverse | CCATCCAATCGGTAGTAGCG |
| UCP1 | Forward | CGACTCAGTCCAAGAGTACTTCTCTT |
|  | Reverse | GCCGGCTGAGATCTTGTTTC |
| CIDEA | Forward | TCTGCAATCCCATGAATGTC |
|  | Reverse | CAGTGATTTAAGAGACGCGG |
| COX8B | Forward | TGTGGGGATCTCAGCCATAGT |
|  | Reverse | AGTGGGCTAAGACCCATCCTG |
| PGC1A | Forward | TATGGAGTGACATAGAGTGTGCT |
|  | Reverse | CCACTTCAATCCACCCAGAAAG |
| DIO2 | Forward | TGCGCTGTGTCTGGAACAG |
|  | Reverse | CTGGAATTCGGAGCATCTTCA |
| PERK | Forward | GAGAAGACGCCCATCACCC |
|  | Reverse | TTAGCTCGCCACACACGCT |
| CHOP | Forward | CTGGAAGCCTGGTATGAGGAT |
|  | Reverse | CAGGGTCAAGAGTAGTGAAGGT |
| ATF4 | Forward | CCTTCGACCAGTCGGGTTTG |
|  | Reverse | CTGTCCCGGAAAAGGCATCC |
| XBP1S | Forward | CTGAGTCCGAATCAGGTGCAG |
|  | Reverse | GTCCATGGGAAGATGTTCTGG |
| ATG6 | Forward | TGAACTTCGAGGATGGGTTC |
|  | Reverse | GAATTTGAGCCCTGTTCCAG |
| NAMPT | Forward | TGCCGTGAAAAGAAGACAGA |
|  | Reverse | ACTTCTTTGGCCTCCTGGAT |
| NMANT1 | Forward | GCTGGCCAAGGACTATATGC |
|  | Reverse | GAGCCCTTTCTTCTTGTACGC |
| NMANT2 | Forward | CCATGACTCCTACGGAAAACA |
|  | Reverse | CAGTCGGAATTCTGGACAGC |
| NMANT3 | Forward | CCAGAGACCACCTACACCAAA |
|  | Reverse | CCAGGTCTTTCTTCCCATAGC |
| NAPRT | Forward | GCCCACTCCTTTGTCACTTC |
|  | Reverse | GTAGAGGCACACACGCTTCA |
| NADSYN1 | Forward | TTCTGGACTCTCCGGTCACT |
|  | Reverse | CATCTTGGGTCTGATGAGCA |
